# Brain morphology network alterations in adolescents with autism spectrum disorder: a sex-stratified study

**DOI:** 10.1101/2025.08.28.672884

**Authors:** Nooshin Safari, Isaac Levy, Aiying Zhang, Chirag Agarwal, Jesus M. Cortes, Kevin A. Pelphrey, John Darrell Van Horn, Javier Rasero

**Affiliations:** School of Data Science, University of Virginia, Charlottesville, VA, USA; Computational Neuroimaging Laboratory, BioBizkaia Health Research Institute, Barakaldo, Bizkaia, Spain; Ikerbasque, Basque Foundation for Science, Bilbao, Spain; Department of Cell Biology and Histology, University of the Basque Country (UPV/EHU), Leioa, Bizkaia, Spain; Department of Neurology, University of Virginia, Charlottesville, VA, USA; Department of Psychology, University of Virginia, Charlottesville, VA, USA

**Keywords:** autism spectrum disorder, sex differences, morphological networks, adolescent neuroimaging, cortical features, comorbidity, multivariate analysis

## Abstract

Neuroimaging studies based on altered functional and structural networks have contributed to better characterizing males and females with Autism Spectrum Disorder (ASD), advancing our understanding of the male prevalence in diagnosis. However, much less is known about how brain networks are altered from a morphological perspective, and whether these alterations may help explain sex-related characteristics in ASD. Here, we used structural MRI from a sex- and diagnosis-balanced sample of 337 individuals in typical neurodevelopmental ages (8–18 years) from the Autism Center of Excellence to elucidate sex-specific alterations in morphology-based connectivity, calculated as the similarity between region-wise multivariate morphological signatures. Network-based statistics showed that ASD males had significantly increased connectivity involving the fusiform gyrus, medial orbitofrontal, entorhinal, and parahippocampal cortices. This network profile was further linked to a core social–communication trait within the autistic group. In females with ASD, increased connectivity was found in a subnetwork primarily implicating the entorhinal cortex, followed by the inferior parietal lobule and lateral occipital cortex. In contrast to males, females’ fusiform gyrus showed decreased connectivity with the superior temporal sulcus. No overlap between male- and female-specific profiles was found. Together, these findings offer new insights into the neurobiology underlying sex differences in autism.

## INTRODUCTION

Autism Spectrum Disorder (ASD) is a neurodevelopmental condition defined by persistent impairments in social communication alongside restricted and repetitive behaviors, currently affecting approximately one in 36 children in the United States, with a male-to-female diagnosis ratio ranging from 3:1 to 4:1 [1,2]. The basis of this sex bias is still under debate. Some theories suggest that ASD is characterized by a cognitive trait dominated by systemizing rather than empathizing. Systemizing is more commonly observed in males, which may explain the higher prevalence of autism in this sex. Under this view, females are at the opposite end of the cognitive profile due to their generally stronger ability for empathizing [3]. Another compelling theory is the Female Protective Effect, which posits that females are more resilient to developing autism and require a greater etiological load to reach diagnosis [4]. If these theories are true, they should be at least partially reflected in the brain through sex-specific neural correlates. To examine this, sex should be treated as a stratifying variable rather than a nuisance covariate [5,6].

Much of our understanding of the neurobiology underlying sex-specificity in autism has emerged from neuroimaging efforts evaluating disruptions in brain circuitry. From a functional perspective, females with ASD typically show a trend toward network-wide hyperconnectivity relative to sex-matched typically developing controls (TDC), while males tend toward hypoconnectivity [7]. At a more local level, these patterns can be more nuanced or even reversed [8]. In particular, functional disruptions in females with ASD are predominantly found in somatomotor regions, while males exhibit alterations in both the default mode and somatomotor networks—supporting the so-called Gender Incoherence model [9]. Additional differences in visual, language, and basal ganglia connectivity have also been reported among autistic females [10]. Similar sex disparities have been found in structural connectivity. In preschool-aged children, males with ASD show reduced callosal projections to the orbitofrontal cortex, whereas females exhibit reductions targeting anterior frontal areas, along with elevated diffusivity metrics [11]. During adolescence, females with ASD—but not males—show reduced fiber density and cross-section in callosal pathways [12]. Region-specific reductions in connectivity involving temporal, parietal, and medial posterior cortices have also been reported in females, with no corresponding effects in males [13].

Beyond function and structure, brain morphology has also contributed to further characterizing sex-specific differences in autism. Recent findings indicate that autistic girls exhibit thicker cortices in early development, followed by accelerated thinning relative to autistic boys, suggesting a sex-linked protective mechanism [14]. Overall, morphological features (e.g., cortical thickness, surface area, sulcal depth, mean curvature, and volume) are thought to reflect the cumulative outcome of developmental and genetic processes, positioning them as important markers for disentangling heterogeneity in ASD [15]. Yet, most of this work has focused on region-by-region analyses, typically examining features independently across predefined brain areas, which may be insufficient to capture the spatial interdependencies in brain morphology that sustain neurodevelopment [16].

One promising approach to overcome this limitation is to also adopt a connectivity perspective on brain morphology and examine the similarity in structural features (e.g., cortical thickness, surface area) between brain regions. Most efforts in this direction have employed structural covariance networks, which model connectivity as inter-regional covariation in a single morphometric property across individuals [17,18]. In the context of autism, studies have reported widespread alterations in connectivity based on region-wise volumetric covariation, with some of these patterns linked to socio-cognitive differences [19,20]. A recent study further identified significant sex-by-diagnosis interactions in this type of networks within major brain systems, showing that ASD males had increased covariance relative to typically developing males, while ASD females displayed reduced covariance compared to female controls [21]. Despite these findings, whole-brain, sex-specific alterations in morphology-based connectivity in autism remain largely underexplored. Moreover, covariation network approaches are typically limited to a single morphological property and may overlook the benefits of integrating multiple features into a unified framework to provide a more comprehensive description of group and individual differences in cortical organization [22,23].

To fill these gaps, the present study set out to characterize sex-specific connectivity patterns based on multivariate morphological information in individuals with autism during typical developmental ages. Following similar observations in functional and structural connectivity, we hypothesized that males and females with ASD would show distinct morphological network profiles. To this end, we leveraged a large-scale, multi-site dataset comprising 337 youth participants (aged 8–18 years), balanced by sex (162 females) and diagnosis (181 with ASD). Adopting a recently developed framework for robustly estimating similarity networks from T1-weighted images [24], morphology-based connectivity was computed from multivariate patterns of cortical features, including thickness, mean curvature, surface area, sulcal depth, and gray matter volume. Clusters of statistically disrupted connections in ASD relative to TDC were then identified separately in males and females, and their associations with behavioral and comorbidity traits were further evaluated.

## MATERIALS AND METHODS

### Participants

337 individuals from Wave 1 of NIH Autism Centers of Excellence (ACE) Network were selected for this study after quality control assurance, involving visual inspection of segmentation output and exclusion of scans with structural preprocessing failures (see [25] for more details on the ACE network). The study sample included 181 subjects with ASD (Age: 12.73 ± 2.92 years, 84 female, 136 identifying as White, 8 identifying as Black, 4 identifying as Asian, 28 identifying as multiracial, 2 reporting other racial identities, 3 reporting no racial identity, 147 identifying as not Hispanic, 31 identifying as Hispanic, and 3 reporting no ethnic identity) and 156 with TDC (Age: 13.20 ± 3.01 years, 78 female, 115 identifying as White, 13 identifying as Black, 11 identifying as Asian, 16 identifying as multiracial, 1 reporting no racial identity, 117 identifying as not Hispanic, 36 identifying as Hispanic, and 3 reporting no ethnic identity), who were collected across four sites: Harvard University, Seattle Children’s/University of Washington, University of California Los Angeles, and Yale University (Figure 1A). Additionally, several scores were used to assess the association of altered networks with behavioral and comorbidity traits within individuals with autism. Specifically, we extracted scores for communication, social interaction, and restricted, repetitive, and stereotyped behaviors from the Autism Diagnostic Interview–Revised (ADI-R); social affect and restricted and repetitive behaviors from the Autism Diagnostic Observation Schedule—2nd Edition (ADOS-2); and subscale scores for anxiety, affective problems, attention deficit/hyperactivity, oppositional defiant, conduct, and internalizing problems from the Child Behavior Checklist (CBCL). Descriptive statistics for our sample can be found in Table 1.

**Figure 1.**
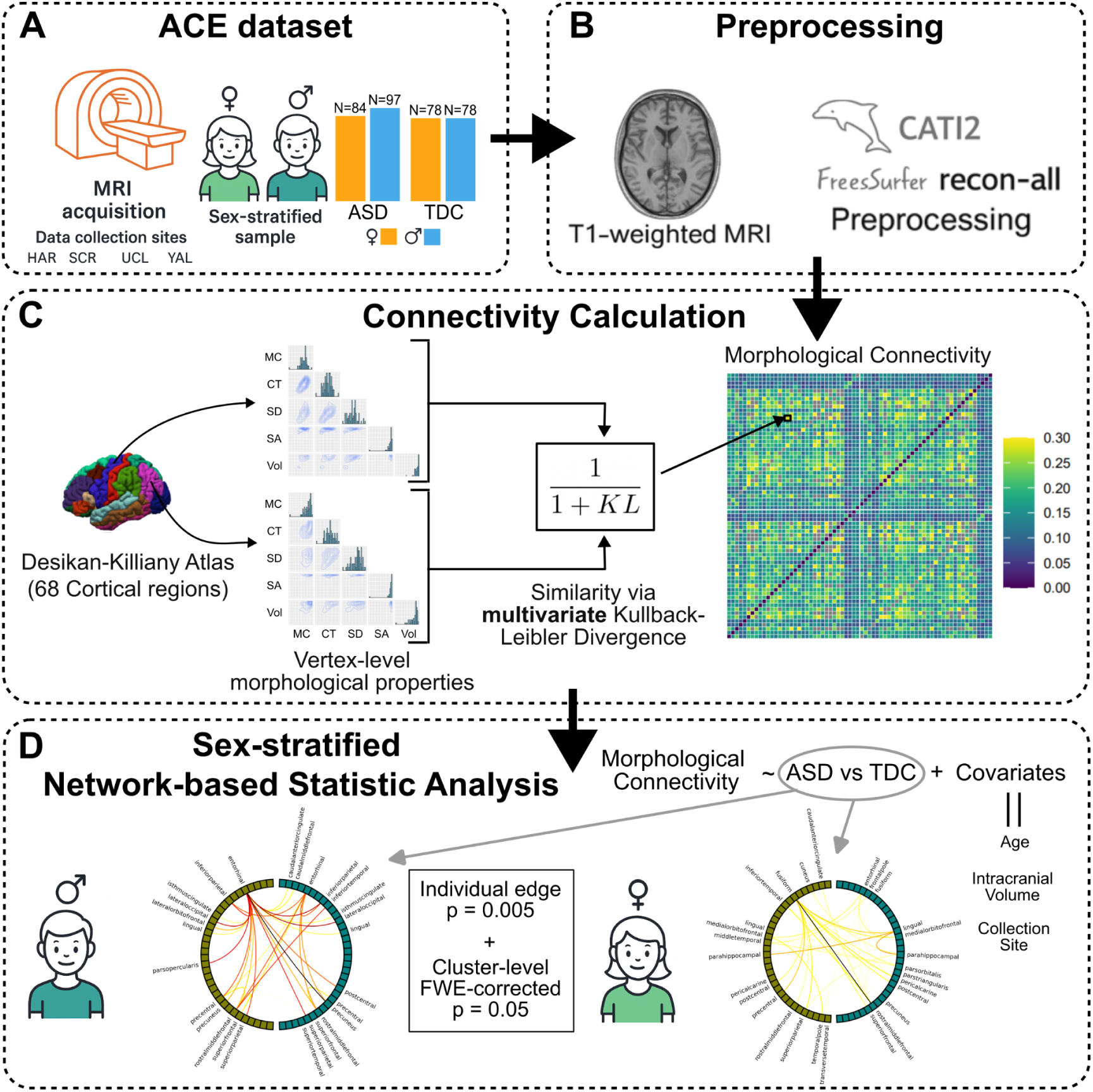
Methodological Workflow. **A)** We analyzed a sex- and diagnosis-balanced sample of 337 individuals aged 8–18 years. **B)** T1w images were preprocessed using FreeSurfer and CAT12 for voxel-based morphometry. **C)** From the preprocessed data, we extracted regional distributions of multivariate morphological features (mean curvature, cortical thickness, sulcus depth, surface area, and volume) defined by the Desikan–Killiany cortical atlas. Inter-regional connectivity was then estimated using the Kullback–Leibler divergence between these multivariate distributions. **D)** Finally, connectivity matrices were regressed onto group labels while controlling for covariates (age, intracranial volume, and collection site), adopting a network-based statistic approach. Analyses were performed separately by sex.

**Table 1.** Demographic, behavioral and comorbidity data stratified by sex and diagnostic group. Values represent means and 95% confidence intervals. Dashes (—) indicate unavailable scores for TDC participants. HAR: Harvard University; SCR: Seattle Children’s/University of Washington; ICV: intracranial volume

|  | Male |  | Female |  |
| --- | --- | --- | --- | --- |
| Variable | ASD | TDC | ASD | TDC |
| N | 97 | 78 | 84 | 78 |
| Age (months) | 152.36<br>[145.02–159.69] | 161.02<br>[153.47–168.57] | 153.34<br>[146.03–160.65] | 155.82<br>[147.10–164.55] |
| Site<br>(HAR/SCR/UCL/YAL) | 24/28/30/15 | 11/23/26/18 | 11/25/25/23 | 13/25/23/17 |
| ICV ($10^6 \times \text{mm}^3$) | 1.63<br>[1.60–1.66] | 1.64<br>[1.61–1.67] | 1.47<br>[1.44–1.50] | 1.48<br>[1.45–1.51] |
| ADI-R Social | 19.38<br>[18.28–20.49] | — | 18.11<br>[16.74–19.48] | — |
| ADI-R Communication | 16.54<br>[15.67–17.42] | — | 15.38<br>[14.37–16.40] | — |
| ADI-R Behavioral | 6.63<br>[6.12–7.14] | — | 5.70<br>[5.13–6.28] | — |
| ADOS-2 Social Affect | 9.52<br>[8.75–10.28] | — | 8.06<br>[7.40–8.72] | — |
| ADOS-2 Communication | 2.63<br>[2.29–2.97] | — | 2.54<br>[2.15–2.93] | — |
| CBCL Affective | 61.07<br>[59.28–62.85] | 51.60<br>[50.86–52.33] | 64.19<br>[61.93–66.46] | 52.95<br>[51.87–54.02] |
| CBCL Anxiety | 61.74<br>[60.13–63.36] | 51.90<br>[51.06–52.73] | 62.49<br>[60.56–64.43] | 52.04<br>[51.14–52.94] |
| CBCL ADHD | 61.92<br>[60.37–63.48] | 50.51<br>[49.00–52.01] | 63.38<br>[61.47–65.28] | 51.18<br>[50.59–51.77] |
| CBCL<br>Oppositional | 58.65<br>[57.04–60.26] | 51.62<br>[50.96–52.28] | 58.38<br>[56.67–60.09] | 51.53<br>[50.94–52.13] |
| CBCL Conduct | 56.17<br>[54.70–57.63] | 50.81<br>[50.32–51.29] | 57.33<br>[55.56–59.10] | 50.86<br>[50.38–51.35] |
| CBCL<br>Internalizing | 60.87<br>[59.15–62.58] | 43.81<br>[41.66–45.95] | 63.04<br>[60.48–65.60] | 46.47<br>[44.31–48.63] |

### MRI Acquisition and Preprocessing

Structural images were acquired using a high-resolution T1-weighted MPRAGE sequence (FOV: 176×256×256; voxel size: 1×1×1 mm^3^). Preprocessing of these images was performed with FreeSurfer’s recon-all pipeline (v6.0), which involves cortical surface reconstruction and subcortical segmentation. Additionally, gray matter volume maps were generated using a standard voxel-based morphometry pipeline implemented in CAT12 [26], and transformed into the FreeSurfer subject space (Figure 1B). These maps enabled the extraction of voxel-wise gray matter distributions for cortical and subcortical regions, needed to compute whole-brain connectivity matrices (see next section). This included spatial normalization, modulation, and smoothing of gray matter probability maps. On average, these images exhibit satisfactory quality (78. 07 ± 5. 28), as quantified using the weighted Image Quality Rating (IQR) metric [26].

Of note, motion-related quality assurance metrics (e.g., framewise displacement) are not applicable here, as our morphology-based connectivity analysis relies on single high-resolution structural images rather than time-series data (e.g., fMRI) where motion effects are more pronounced.

### Morphology-Based Connectivity

Morphology-based connectivity matrices were constructed for all subjects using the Morphometric INverse Divergence framework [24], which estimates pairwise similarity between brain regions based on the symmetrized Kullback–Leibler (KL) divergence of vertex-wise morphometric feature distributions (Figure 1C). Main analyses concentrated on cortical networks, where multiple morphological properties could be defined. Additional analyses were performed on whole-brain networks, but these were based solely on volumetric information, since this was the only morphological measure available for subcortical regions.

Specifically, to build the connectivity matrices at the cortical level, the vertex-level outputs for thickness, mean curvature, surface area, sulcal depth, and gray matter volume provided by FreeSurfer were extracted for each region of the Desikan–Killiany atlas (68 regions of interest, ROI; 34 per hemisphere), i.e., within each ROI, a distribution of vertex-wise values for each of the aforementioned morphological properties was obtained. These distributions were then modeled with kernel density estimation and jointly used to compute the similarity between pairs of regions via a symmetrized multivariate KL divergence. This resulted in a 68 x 68 connectivity matrix for each individual, representing the similarity between regions based on their multivariate morphological signatures.

For the whole-brain connectivity matrices, 14 subcortical structures obtained from FreeSurfer’s *aseg.mgz* segmentation file [27] were added to the aforementioned cortical regions from the Desikan–Killiany atlas. These subcortical regions consisted of the bilateral thalamus, caudate, putamen, pallidum, hippocampus, amygdala, and nucleus accumbens. An 82 x 82 connectivity matrix was then built for each individual as the univariate KL divergence between region-wise gray matter volume distributions, extracted from the voxel values within each ROI after the voxel-based morphometry pipeline.

### Sex-Specific Network Alterations

To identify statistically significant differences in morphology-based connectivity, we applied the Network-Based Statistics (NBS) approach [28], which is analogous to cluster-wise thresholding of statistical brain maps but adapted for network data (Figure 1D). For each edge in the connectivity matrix, a test statistic was first computed, followed by an individual threshold to identify sub-networks of suprathreshold connected edges. The significance of each sub-network was then assessed using permutation testing while controlling the family-wise error rate. A two-sided linear regression model was employed to calculate the test statistic at each edge, therefore allowing the existence of significant clusters of both increased and decreased connectivity in ASD relative to TDC. For interpretability, the resulting edges were subsequently separated based on the sign of the observed test statistic. This analysis was implemented using the *lm_nbr* function from the NBR R package [29].

The current study concentrated on **sex-specific** network alterations. As a result, NBS was applied to the connectivity matrices within each sex subpopulation (male and female) with the diagnosis variable (ASD vs. TDC) as the main effect, while controlling for covariation in age, intracranial volume, and collection site. By including intracranial volume as a covariate, we ensured that group effects reflect true connectivity differences rather than global volumetric scaling. Testing a traditional sex × diagnosis interaction effect was considered. However, since females and males typically differ in overall cortical morphology—including well-established differences in intracranial volume (see Table 1)—interaction-based models risked conflating diagnosis-related effects with sex-specific variance structure. Therefore, we adopted a more conservative and interpretable approach by conducting analyses separately within each sex. Main results were reported for NBS with 5000 total permutations, an individual edge threshold *p* = 0. 005 and family-wise error correction (*p* _*FWE*_ = 0. 05). In NBS, the individual edge threshold is a hyperparameter that determines the potential size of clusters. We chose *p* = 0. 005 since it provided a compromise in cluster size, sufficient for discovery while preserving statistical power [28]. To assess robustness, we also tested the sensitivity of the findings to a more liberal (*p* = 0. 01) and stringent threshold (*p* = 0. 001).

### Importance of Morphological Properties

The relative importance of each morphological property in distinguishing individuals with autism from typically developing controls was assessed by combining the strength of each sub-network identified by NBS with a leave-one-feature-out procedure within an ROC curve analysis framework. Strength was calculated for each individual’s connectivity matrix as the sum of values corresponding to the altered sub-networks. This metric reflects the overall magnitude of morphological similarity within the significant sub-networks and provides a compact summary of network-level disruption. The leave-one-feature-out procedure removed a given morphological property prior to constructing the connectivity matrices via multivariate KL divergence, thereby changing the strength of the sub-networks too. The impact on classification performance was then quantified by examining the change in the area under the ROC curve (AUC) using strength as the discriminating variable. This procedure was repeated for each morphological property individually.

### Association with Behavior and Comorbidity

Within the autistic group, we applied Partial Least Squares (PLS) regression to test the association between the significantly altered connectivity edges identified by NBS and multivariate behavioral information, comprising the social, communication, and behavioral subscores from ADI-R, as well as the behavioral and social affect scores from ADOS-2. PLS aims to find latent components such that a multivariate dataset *X* (connectivity values of significantly altered edges by NBS) maximally covaries with another multivariate dataset *Y* (here, multivariate behavioral information). Null distributions for each latent component were generated by repeatedly running PLS with the observation-wise permutation of *Y* (5000 permutations), and statistical significance was claimed at the 0.05 threshold after FDR correction. Furthermore, contributions of individual features in *X* and *Y* to each significant latent component were assessed by inspecting the corresponding loading profiles, focusing on those features whose loadings significantly deviated from zero at a 95% confidence interval calculated by bootstrapping (5000 iterations). This analysis was implemented using the *pyls* Python package (https://pyls.readthedocs.io/en/latest). To maintain consistency with the covariates used in the primary NBS analysis, the effects of age, site, and intracranial volume were regressed out from both connectivity- and behavior-based datasets before the PLS regression.

The same approach was adopted to assess the relationship between the significantly altered connectivity edges and comorbidity symptoms, consisting of affective problems, anxiety problems, attention deficit/hyperactivity, oppositional defiant problems, conduct problems, and internalizing problems scores in CBCL.

## RESULTS

### ASD Males vs. TDC Males

Group-level network analysis identified a single cluster of 35 significantly altered connections (*p* _*FWE*_ = 0. 037). As shown in Figure 2A, most of these corresponded to increased morphology-based connectivity in ASD relative to TDC, with a pattern prominently involving the left hemisphere fusiform and medial orbital frontal gyri, as well as the right hemisphere entorhinal cortex and lingual gyrus. This cluster also included a single connection showing decreased connectivity in ASD compared to TDC, linking the left middle temporal gyrus and the right inferior temporal gyrus. This analysis was performed using the individual edge threshold *p* = 0. 005 and 5000 permutations for null modeling. Additional profiles with other individual edge thresholds in NBS were also reported (see Supplementary Fig. 1). Remarkably, the fusiform continued to exhibit significantly decreased connectivity even under the most stringent threshold (*p* = 0. 001). No significantly altered networks at the whole-brain level using volumetric information only were found at any of the thresholds.

**Figure 2:**
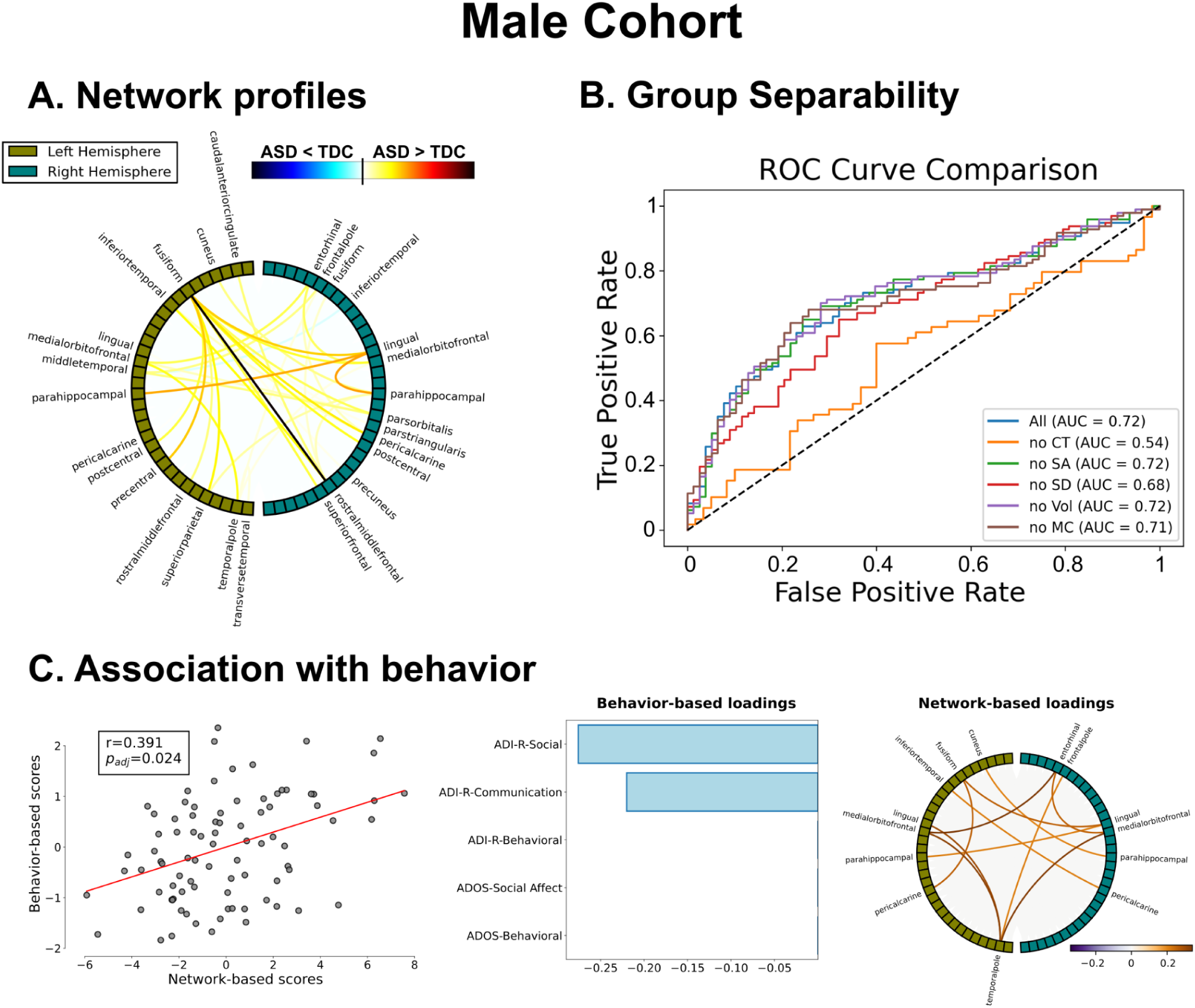
Main results in the male cohort. **A)** Circular plots visualizing clusters of significant inter-regional connections. Nodes represent cortical regions, with olive and teal indicating the left and right hemispheres, respectively. Blue (red) colors indicate increased (decreased) connectivity in TDC relative to ASD. Darker colors denote stronger group differences. **B)** ROC curves and area under the curve (AUC) values showing performance in distinguishing ASD from TDC using the strength of network profiles in **A**. Each curve represents the scenario in which one morphological property is omitted in the computation of the connectivity matrices (CT: cortical thickness, SA: surface area, SD: sulcus depth, Vol: volume, MC: mean curvature). **C)** Association between morphology-based connectivity and behavior in ASD provided by a significant latent component using Partial Least Squares regression (left panel), and their corresponding loadings (middle and right panel), focusing only on those contributing factors whose weights are away from zero at a 95% confidence level.

The most influential morphological property for distinguishing between controls and individuals with autism within this cluster appeared to be cortical thickness, with the AUC dropping from 0.72 (when all features were included) to 0.54 when this feature was removed (see Figure 2B). This was followed by sulcal depth (AUC = 0.68) and mean curvature, although in the latter case the reduction in group separability was minimal (AUC = 0.71). In contrast, surface area and volumetric properties did not appear to be individually necessary, as removing either of them resulted in slightly improved group separability (AUC = 0.72 in both cases).

We found a latent component by PLS (r=0.391, *p_adj_* = 0.024) in which altered morphological connections significantly associate with behavior within the autistic group (left panel in Figure 2C). This association was primarily driven by negative loadings of social and communication scores from ADI-R (middle panel in Figure 2C) and positive loadings of the left hemisphere fusiform and temporal pole regions, as well as the right hemisphere entorhinal and lingual cortices (right panel in Figure 2C), indicating a negative correlation between ASD-relevant behavior and altered morphological connectivity profiles.

No significant latent components between altered sub-networks and comorbidity information were found after FDR correction.

### ASD Females vs. TDC Females

Within the female subgroup, a single cluster comprising 41 significantly altered morphology-based connections (*p* _*FWE*_ = 0. 028) was identified. As displayed in Figure 3A, compared to TDC, patterns of increased connectivity in the ASD group were centered around a prominent hub in the bilateral entorhinal cortex, which then extended to the bilateral inferior parietal lobule, lateral occipital cortex, and pre- and postcentral gyri, all regions implicated in visuospatial integration, somatosensory encoding, and mnemonic retrieval. In contrast, decreased connectivity, although more scarce, prominently involved the bank of the superior temporal sulcus engaging with a few other regions, among which the fusiform gyrus was found. All these alterations remained consistent—yet spatially circumscribed, more notoriously to the left hemisphere entorhinal cortex—at the more stringent individual edge threshold in NBS (*p* = 0. 001). At the more liberal *p* = 0. 01 level, the pattern expanded to include bilateral precentral extensions and deeper parietal-occipital couplings (see Supplementary Fig. 2), suggesting a core-periphery organization sensitive to statistical stringency. Similar to males, no significantly altered networks were found at any of the thresholds at the whole-brain level using only volumetric information.

**Figure 3:**
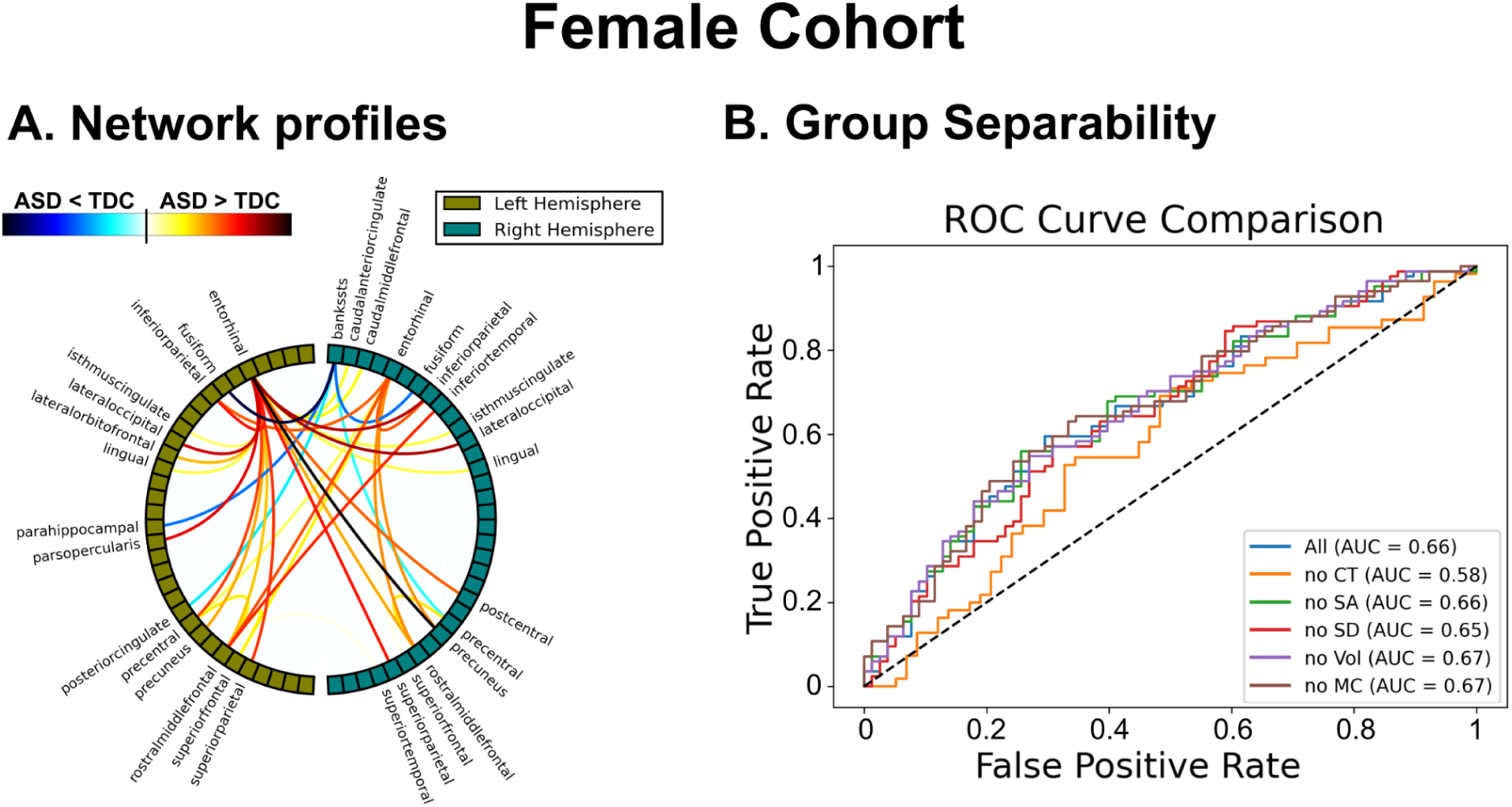
Main results in the female cohort. **A)** Circular plots visualizing clusters of significant inter-regional connections. Nodes represent cortical regions, with olive and teal indicating the left and right hemispheres, respectively. Blue (red) colors indicate increased (decreased) connectivity in TDC relative to ASD. **B)** ROC curves and area under the curve (AUC) values showing performance in distinguishing ASD from TDC using the strength of network profiles in **A**. Each curve represents the scenario in which one morphological property is omitted in the computation of the connectivity matrices (CT: cortical thickness, SA: surface area, SD: sulcus depth, Vol: volume, MC: mean curvature).

In terms of group separability, a similar pattern to that observed in the male cohort was found, to a large extent (see Figure 3B). Using all morphological features together yielded an AUC of 0.66. Removing cortical thickness led to the largest reduction in group separability (AUC = 0.58), confirming its key role. In contrast, removing surface area had no effect (AUC = 0.66), while sulcal depth had only a minimal impact in this case (AUC = 0.65). Volumetric properties and mean curvature individually did not appear to be essential, as their removal slightly increased the AUC to 0.67 in both cases.

No significant latent components linking altered morphological network profiles with either behavioral or comorbidity information in females with autism were found after FDR correction.

### Network Profiles Comparison

We conducted an overlap analysis to assess the distinctiveness of the network profiles identified across sexes. For each group comparison (i.e., ASD males vs. TDC males, ASD females vs. TDC females), we first constructed a binary adjacency matrix in which an entry was set to 1 if the corresponding edge was found to be significantly altered by NBS. We then computed the Jaccard index (*J*) defined as the number of intersecting positive cases divided by the size of their union, i.e., 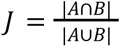 At the individual edge threshold of *p* = 0. 005 and cluster-level family-wise error correction (*p* _*FWE*_ = 0. 05), this yielded a Jaccard index of *J* = 0 between the male and female network profiles, indicating no overlapping edges across the sex-specific contrasts.

To further evaluate whether these profiles reflected true diagnostic group effects rather than normative sex differences, we applied the same NBS setup of our main analysis (individual edge threshold *p* = 0. 005, cluster-level family-wise error correction *p* _*FWE*_ = 0. 05, 5000 permutations, and identical covariates) to compare TDC males with TDC females. This analysis revealed a cluster of 48 significantly altered connections, which are depicted by directionality in Supplementary Fig. 3. When overlapping this normative sex-difference network with the diagnostic-related profiles, we found a Jaccard index of *J* = 0 for ASD males vs. TDC males, and *J* = 0. 01 for ASD females vs. TDC females. In the latter case, only a single overlapping connection was found, linking the left rostral middle frontal gyrus and the right superior temporal gyrus. These minimal overlaps suggest that the reported sex-specific morphological network profiles arise from diagnosis-related alterations rather than normative sex differences.

## DISCUSSION

This study aimed to delineate sex-specific network profiles based on altered brain morphology-based connectivity in adolescents with autism. To accomplish this, we leveraged a large-scale, multi-study dataset comprising 337 individuals, balanced by sex and diagnostic group. For each participant’s T1-weighted image, we extracted vertex-level distributions of cortical morphometric properties across 68 brain regions, including cortical thickness, mean curvature, surface area, sulcal depth, and gray matter volume. We then estimated inter-regional connectivity by quantifying the similarity between these morphometric distributions using a recently proposed approach based on multivariate KL divergence. By means of a network-based statistical framework, we assessed sex-specific alterations in connectivity between ASD and TDC individuals and found distinct profiles across males and females. Specifically, males with autism showed significantly heightened levels of connectivity in regions prominently involving the fusiform and lingual gyri, anterior cingulate cortex, orbitofrontal cortex, and parietal association cortices, whereas for females, statistical results centered on elevated connectivity in the entorhinal cortex, inferior parietal lobule, and lateral occipital cortex, as well as a decreased connectivity between the fusiform gyrus and the bank of superior temporal sulcus. Our results suggest that morphology and connectivity can further characterize autism and, importantly, provide novel brain-based insights into the sex-specific differences commonly observed in individuals with this neurodiverse condition.

Our results add to the growing body of evidence reinforcing a network-level perspective for better characterizing sex differences in ASD. In males, we found increased connectivity in regions belonging to the default mode, salience, and frontoparietal networks. One of the central hubs we identified is the fusiform gyrus, a region implicated in social cognition that has been consistently linked to autism through findings of reduced activation during face processing and structural abnormalities associated with social perceptual deficits [30–32]. Other key regions included the parahippocampal gyrus (linked to memory) and the postcentral gyrus (involved in sensorimotor processing). These findings align with theories of network-level developmental plasticity, in which increased long-range integration may reflect delayed synaptic pruning or compensatory adaptations to altered sensory or cognitive demands, as supported by studies using graph-theoretical metrics [33,34]. Importantly, these areas have also been found to be hyperconnected in fMRI studies using male-predominant cohorts [35–37] and, as we previously demonstrated, contribute to the functional connectivity–based heterogeneity observed in autism [38]. Furthermore, they have also been identified as part of a disrupted structural architecture in ASD, exhibiting reduced fractional anisotropy and aberrant streamline coherence in white matter tracts traversing the cingulum bundle, forceps minor, and superior longitudinal fasciculus [39–41]. Taken together, our findings point to a coherent pattern of multimodal dysconnectivity in males with ASD, centered on frontal, temporal, and occipital integration hubs as key components of ASD neuropathophysiology.

On the other hand, in females with ASD, network disruptions across both functional and structural modalities also appeared to overlap with key regions identified in our morphology-based connectivity profiles. Specifically, resting-state fMRI studies have revealed disruptions within the default-mode network (heightened entorhinal–posterior cingulate; [42]), altered top-down central executive influence—specifically dorsolateral prefrontal to postcentral pathways [43]— and increased coupling between the default-mode and central executive networks (Hong et al., 2019). From the perspective of the canonical triple-network model [44], these changes suggest a reconfigured network topology in ASD females, which may reflect compensatory upregulation of executive–memory loops to offset local cortical variability. Additionally, sex-specific salience network alterations—such as elevated connectivity with sensorimotor areas reported in early ASD trajectories (infants at familial risk) [45]—may reflect dynamic sensory reweighting and flexible attention gating across internal–external stimulus axes. Further, reduced network flexibility in executive and salience systems (as quantified via multilayer network analyses) correlates with social–communication symptom severity in ASD females [46], raising interpretative nuance around whether increased ‘integration’ denotes an adaptive scaffold or constrained network rigidity. In parallel, DTI studies in ASD females aged 12–30 years have revealed significant reductions in fractional anisotropy within white matter tracts underlying our morphological-based network profiles—including the internal capsule, inferior parietal lobule–lateral occipital cortex association fibers, and posterior corona radiata [47,48]. In sum, these convergent findings appear to support the presence of a multimodal (morphology, function, and structure) hub integration profile that may be specific to the female ASD phenotype.

What, then, do these sex-specific morphological network alterations reveal about the broader neurobiology of autism in light of sex differences? Even though our study did not directly contrast sexes, the involvement of largely distinct regions—and in some cases, with opposite directionality—suggests divergent neural substrates underlying core autistic traits. For example, we found hyperconnectivity of the fusiform gyrus in males but hypoconnectivity in females. Since this region plays an established role in face perception and social cognition [49–51], its increased connectivity in males with autism could mirror a compensatory mechanism or atypical social attention pathways typically found in male-biased ASD studies but at the morphological architecture level. In contrast, the reduced connectivity of the fusiform in females, particularly involving connections with the superior temporal sulcus—another social cognitive hub—may signal a qualitatively different neurodevelopmental trajectory. Interestingly, the entorhinal cortex appears to be more disrupted in females than males. As a central node in memory and spatial navigation [52,53], this finding may hint at alternative circuitries supporting adaptive behavior. Critically, these highlighted regions are unlikely to be attributed to normative sex differences, as our overlap analysis demonstrated.

We assessed the sensitivity in discriminating utility of each morphological property underlying the obtained sex-specific connectivity patterns using a one-leave-feature-out procedure. Across sexes, the largest drops in diagnostic performance occurred when cortical thickness and sulcus depth were excluded, suggesting that these properties play a particularly sensitive role in the discriminative capacity of morphological networks—or at the very least, offer complementary information relative to the other morphological properties. Cortical thickness is a well-established marker known to be altered in autism and associated with symptom severity [15,54–56]. Sulcal depth, by contrast, has been less extensively studied in autism, with findings ranging from no statistical differences [57], to specific localized abnormalities [58], and to associations with early cortical folding processes during neurodevelopment [59]. Our study builds on this body of work by highlighting the relevance of both of these morphological properties from a network-level perspective and across sexes. Conversely, removing either surface area or volumetric information from the connectivity calculations led to slight improvements in discriminating utility, which seems to indicate a potentially redundant or less informative contribution from these properties. This can be partially explained by the strong correlation typically observed between surface area and volume at the cortex-level, so very little or none is gained from their combination. Nevertheless, in our analysis based solely on volumetric information, performed to allow the inclusion of the subcortex, we did not find any significant clusters of altered edges across sexes, reinforcing the limited utility of volumetric features to identify ASD individuals, at least from a network perspective. This appears to contradict prior studies suggesting that brain networks based on volumetric features have high predictive power for classifying ASD individuals [20]. However, this may simply reflect the need for a more spatially distributed signature based on volumetric information, whereas our network-based statistical framework focuses on detecting clusters of significantly altered connections and might therefore be less efficient for classification purposes.

To assess whether morphological network profiles can be related to a multivariate behavioral trait, we examined multivariate associations between connectivity—restricted to those entries previously identified by NBS—and core autism-related behavioral measures within the autistic group. We found a significant brain–behavior association in males with autism, suggesting that their inter-individual differences in morphology-based connectivity may capture aspects of behavioral heterogeneity. In contrast, no brain-behavior association was found in females, which is consistent with the possibility that their ASD-related behavior may be more multifaceted and arise from widespread network disruptions across the brain rather than mapping onto discrete and localized clusters of altered connectivity [36,60,61]. On the other hand, no statistical associations with comorbidity were found, suggesting that the observed morphology-based connectivity alterations are unlikely to manifest general traits of other psychiatric conditions.

Several limitations of our study should be acknowledged. First, our analysis was stratified by sex to mitigate potential confounding arising from sex-specific differences in global morphology. As such, direct comparisons across sexes should be interpreted with caution. Future studies with more balanced designs and carefully controlled variables will be needed to directly test for sex-by-diagnosis interaction effects. Second, our data, drawn from Wave 1 of the ACE network, is cross-sectional and therefore precludes any conclusions about the developmental trajectory of our findings. However, subsequent waves (Wave 2 and 3) of the ACE network include follow-up data, which will enable us to test explicitly for longitudinal effects in future avenues. Third, our findings were derived using a purely statistical inference approach, which, although useful for gaining insight into the morphology-based network architecture characteristic of autism and its sex-specific heterogeneity, does not quantify the utility of these features as biomarkers for diagnostic purposes. Future studies should adopt a thorough machine learning framework with out-of-sample testing to specifically assess this. Finally, our network results are restricted to a single neuroimaging modality, specifically, brain morphology. Hence, their combination with biomarkers from functional and structural connectivity signals may lead to a more comprehensive characterization of the ASD heterogeneity and improved diagnostic accuracy rates [62]. Future studies should also examine the robustness and generalizability of sex-specific network effects across modalities [63,64].

Despite the aforementioned limitations, the present study provides evidence for a brain connectivity profile, derived from multivariate morphological signatures, that is expressed differently across sexes in adolescents with autism. This could contribute to improving the identification of autism in female individuals at younger ages, who are typically underdiagnosed.

## Acknowledgements

NS’s work was supported by the UVA School of Data Science. KAP and JDVH were supported by the National Institute of Mental Health (NIMH) Autism Center of Excellence Network Award (R01 MH100028; PI: KAP).

## Ethics approval and consent to participate

The recruitment of participants for this dataset was approved by the Yale Institutional Review Board, the UCLA Office of the Human Research Protection Program, the Boston Children’s Hospital Institutional Review Board, the USC Office for the Protection of Research Subjects, and the University of Virginia Institutional Review Board for Health Sciences Research. All procedures were conducted in accordance with the Declaration of Helsinki. Minor participants provided verbal assent and were informed they could withdraw at any point during the experiment. Written informed consent was obtained from a parent or legal guardian of each participating child.

## Availability of data and materials

Raw neuroimaging data used in this study can be downloaded from the National Institute of Mental Health Data Archive Data (collection #2021). Source files, analysis scripts and figures are available at https://github.com/RaseroLab/Autism-sex-stratified-morphology-connectivity.

## Authors’ contributions

NS, AZ, CA, JMC, KAP, JDVH and JR contributed to the conceptualization of the project and interpretation of findings. KAP led data collection. NS, IL, and JR performed the analysis. NS and JR drafted the manuscript and prepared the figures. All authors edited the manuscript and approved the final version.

## SUPPLEMENTARY INFORMATION

**Fig S1:**
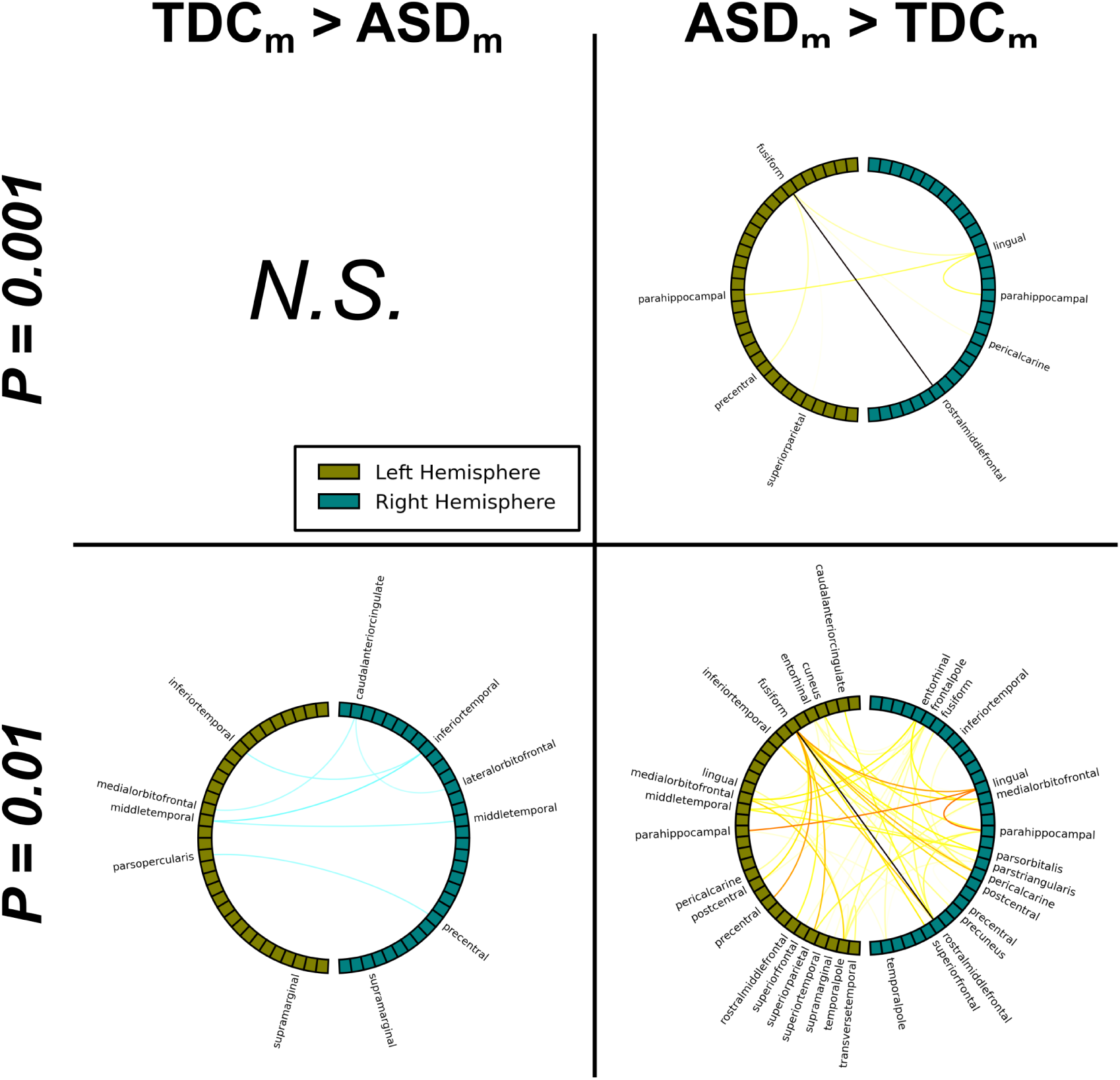
Network profiles with different individual edge thresholds (male cohort). Circular plots visualizing clusters of significant inter-regional connections using a more stringent individual edge threshold (*p* = 0. 001, first row) and a more lenient threshold (*p* = 0. 01, second row), than in the main analysis (*p* = 0. 005). Nodes represent cortical regions, with olive and teal indicating the left and right hemispheres, respectively. Blue colors indicate increased connectivity in the TDC group relative to ASD (left panels), and blue colors the opposite (right panels).

**Fig S2:**
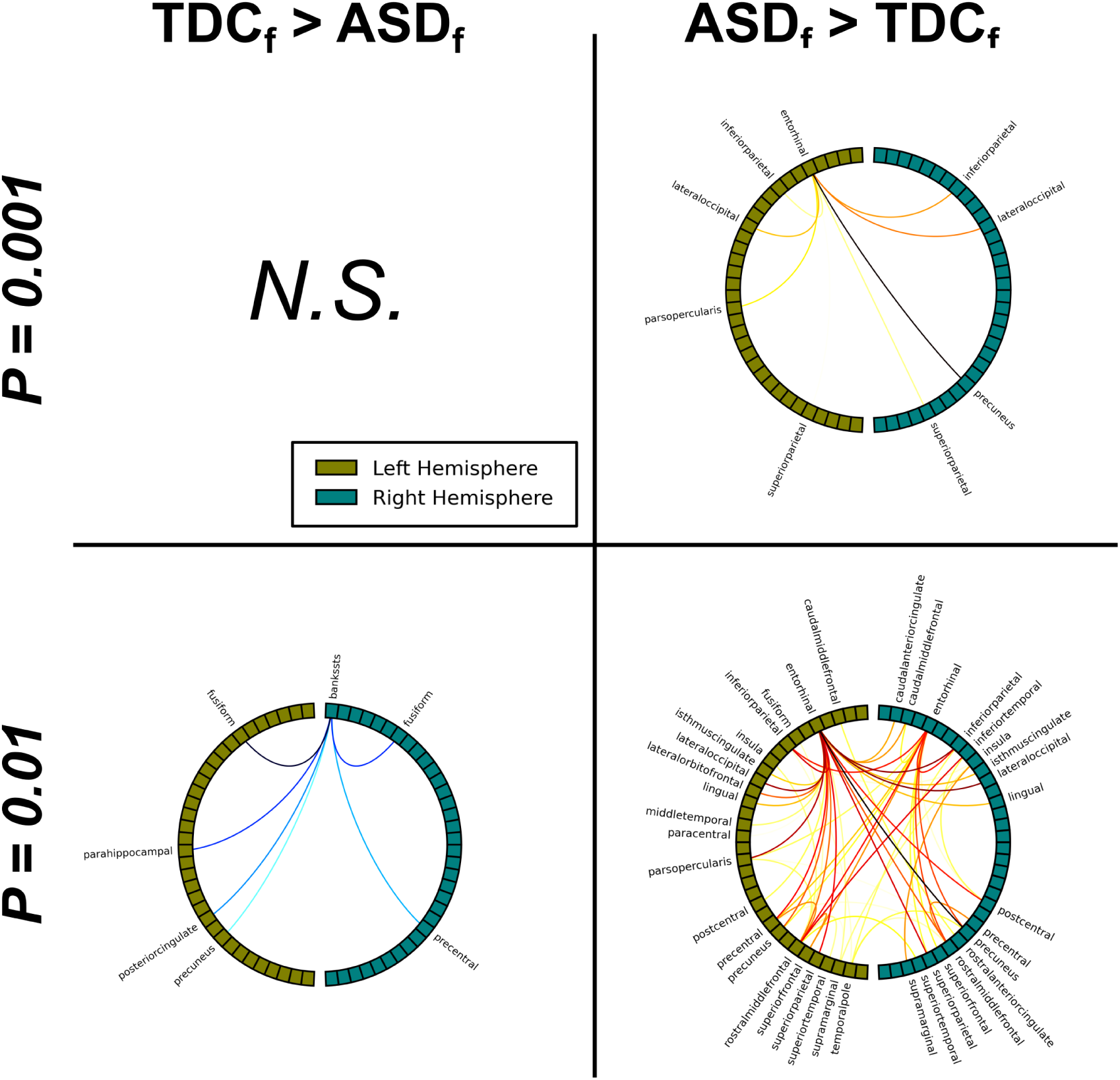
Network profiles with different individual edge thresholds (female cohort). Circular plots visualizing clusters of significant inter-regional connections using a more stringent individual edge threshold (*p* = 0. 001, first row) and a more lenient threshold (*p* = 0. 01, second row), than in the main analysis (*p* = 0. 005). Nodes represent cortical regions, with olive and teal indicating the left and right hemispheres, respectively. Blue colors indicate increased connectivity in the TDC group relative to ASD (left panels), and blue colors the opposite (right panels).

**Fig S3:**
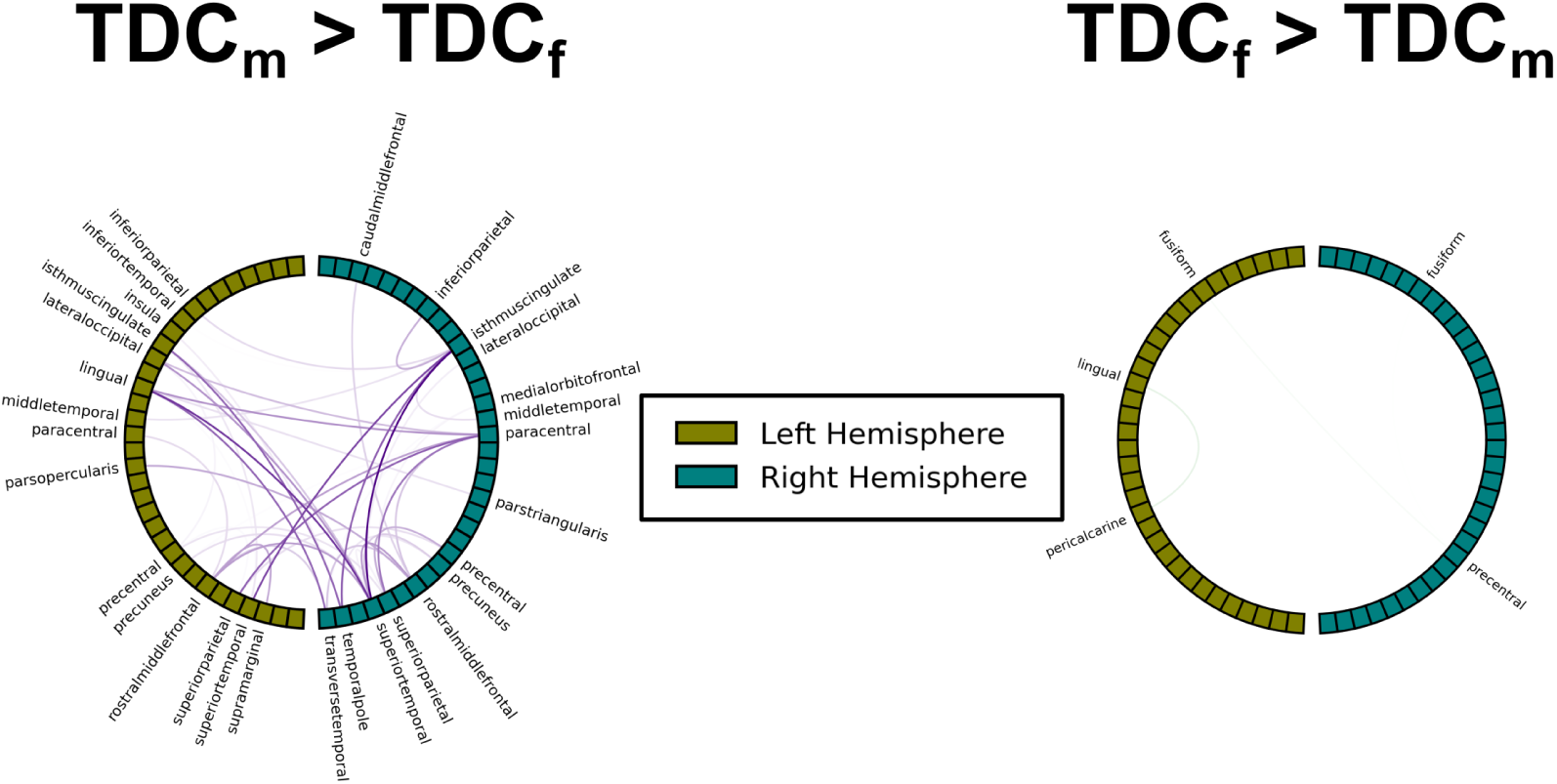
Sex-normative morphological network profiles. Circular plots visualizing clusters of significant inter-regional connections comparing males and females within the TDC group. The same setup as in the main analysis was adopted (individual edge threshold p = 0.005; FWE-corrected p = 0.05; 5,000 permutations). Nodes represent cortical regions, with olive and teal indicating the left and right hemispheres, respectively. Indigo colors indicate increased connectivity in males relative to females, and green the opposite.

